# Mechanical Roles of Vinculin/β-catenin interaction in Adherens Junction

**DOI:** 10.1101/770735

**Authors:** Cristina Bertocchi, Andrea Ravasio, Hui Ting Ong, Yusuke Toyama, Pakorn Kanchanawong

## Abstract

Mechanical force transmission through the adherens junctions (AJs) are highly regulated processes essential for multicellular organization of tissues. AJ proteins such as E-cadherin, α-catenin, and vinculin have been shown to be sensing or bearing mechanical forces being transmitted between the actin cytoskeleton and the intercellular contacts. However, the molecular organization and connectivity of these components remains not well understood. Using a super-resolution microscopy approach, we report that vinculin, once activated, could form a direct structural connection with β-catenin, which can bypass α-catenin, one of the main mechanotransducers in AJs. Direct vinculin/β-catenin interaction is capable of supporting mechanical tension and contributes to the stabilization of the cadherin-catenin complexes. These findings suggest a multi-step model for the force-dependent reinforcement of AJs whereby α-catenin may serve as the initial catalytic activator of vinculin, followed by vinculin translocation to form a direct link between E-cadherin-bound β-catenin and the actin cytoskeleton.

## Introduction

Within epithelial tissues, forces between cells are borne by specialized cell-cell contact structures. The propagation of such forces are essential for developmental morphogenesis, tissue homeostasis, and collective cell migration [1–4], and thus is subject to complex cell-type- and context- specific spatiotemporal regulations. The Adherens Junctions (AJs) are prominent adhesive structures in epithelia that physically couple neighbouring cells via Ca^2+^-dependent homophilic interaction between the extracellular domains of classical cadherins [5–7]. The cadherin receptors in AJs are, in turn, coupled with the actomyosin cytoskeleton via an extensive ensemble of cytoplasmic proteins [8], thereby integrating the mechanobiological processes of cell-cell adhesions with many important cellular pathways[9]. In recent years, the functions of AJs are increasingly better understood across multiple length scales, particularly at the tissues and molecular levels[2]. However, several aspects of AJ functions, especially at the nanoscale regime where AJ proteins are collectively organized into supramolecular assembly, have been largely unexplored until recently [10]. Hence, processes occurring at this length scale, including how mechanical force transmission are mediated and regulated through AJs, remain less well understood.

The ternary complex of epithelial (E-) cadherin, β-catenin, and α-catenin, is widely considered to be the main conduit of mechanical force in AJs [11]. The coupling of actomyosin contractility into intramolecular tension in E-cadherin and α-catenin has been demonstrated using Forster Resonance Energy Transfer (FRET) probes [12–14]. Moreover, cell biological and single-molecule force spectroscopy studies have revealed a tension-dependent conformational changes in α-catenin which gates the recruitment of vinculin, an adaptor protein that reinforces the connection between AJs and the actin cytoskeleton[15, 16]. These proteins were observed by super-resolution microscopy to form a compact cadherin-catenin compartment proximal to the plasma membrane, with vinculin being centrally located between the cadherin-catenin compartment and the actin cytoskeletons [17]. Furthermore, vinculin was found to be conformationally extended upon mechanical and biochemical signals to modulate the AJ nanostructure as well as mechanical tension, consistent with the molecular clutch model [17]. However, given the multi-functional context-dependent roles of many AJ components [18, 19], the wealth of intra-AJ interaction partners[4], and the high-density organization and dynamic plasticity of AJs, these findings could still represented only the most prominent modes of action. Furthermore, the understanding of how these principal AJ components function as an integrated system is still obscure.

Biochemical evidences thus far indicate that the linkage between cadherins and the contractile actin cytoskeleton is indirect [8]. While recent studies have focused largely on the force transmission hierarchy of E-cadherin/β-catenin/α-catenin/vinculin/actin, we note that several alternate pathways may also participate in connecting cadherin with actin, including myosin VI [20], EPLIN [21], α-actinin [22], and afadin [23, 24]. Additionally, β-catenin may interact directly with vinculin [25], which could potentially form an alternate bypass connection from cadherin to the actin cytoskeleton. Collectively, these suggest that in AJs a mixture of different mechanical connectors, rather than a single dominant tether, may couple the cadherin receptors and actomyosin cytoskeletons. However, these alternate interactions have been comparatively under-studied. In this report, we investigated the alternate force transmission pathways in AJs using an integrated approach of super-resolution microscopy, laser nanoscissor, and live-cell microscopy. We found that vinculin, once activated, could form a mechanical connection directly to β-catenin, bypassing α-catenin. However, α-catenin may be required for vinculin activation. These data is suggestive of a multi-step mechanochemical dynamics whereby α-catenin may catalyse vinculin activation in a force-dependent manner, followed by the translocation of vinculin to β-catenin to form a more direct mechanical connection, potentially underpinning AJ reinforcement.

## Results and Discussion

### Super-resolution microscopy reveals an alternate nanoscale organization of vinculin in the absence of α-catenin

In a recent study, we have demonstrated the use of cadherin biomimetic substrate system to facilitate ultra-high resolution imaging of cadherin-based adhesions [17], Fig. 1A. The use of such platform, in conjunction with surface-generated structured illumination microscopy [26, 27], permits sub-20 nm precision analysis along the vertical (z)-axis perpendicular to the plasma membrane. This helped elucidate the nanoscale stratification of proteins into distinct layers, and permit the investigation of conformation changes in proteins such as vinculin [17]. Using this strategy to investigate MDCK (Madin-Derby Canine Kidney) epithelial cells, we investigated the dependence of the nanoscale architecture of E-cadherin-based adhesions on α-catenin, comparing control cells with a stable cell line expressing α-catenin shRNA[28]. As shown in Fig. 1B-C, the fluorophore z-position relative to the substrate surface (z = 0⍰nm) was analysed pixel-wise, with the median value, z_centre_, for adhesion regions of interest (ROIs) used as the representative protein z-position. In both of these conditions, immunostaining showed that endogenous vinculin localizes to the cadherin-based adhesions (Fig. 1E-F). Using fluorescent protein (FP) fusion as probes, we found that E-cadherin (C-terminal GFP fusion) was located at similar z-positions. In contrast, a drastic difference was observed in the z-position of vinculin. As shown in Fig, 1C-D and Supplementary Table S1, in control MDCK, vinculin is positioned proximal to the cadherin-catenin compartment (z <60 nm), but upon α-catenin depletion, vinculin localizes to a much higher z-position (z>80 nm), coinciding with the actomyosin compartment, probably via the association with actin or actin-regulatory proteins such as VASP or α-actinin [29, 30]. As we have shown earlier [17], we found that the central position of vinculin is restored upon re-expression of α-catenin indicative of the role of α-catenin in the central placement of vinculin.

**Figure 1.**
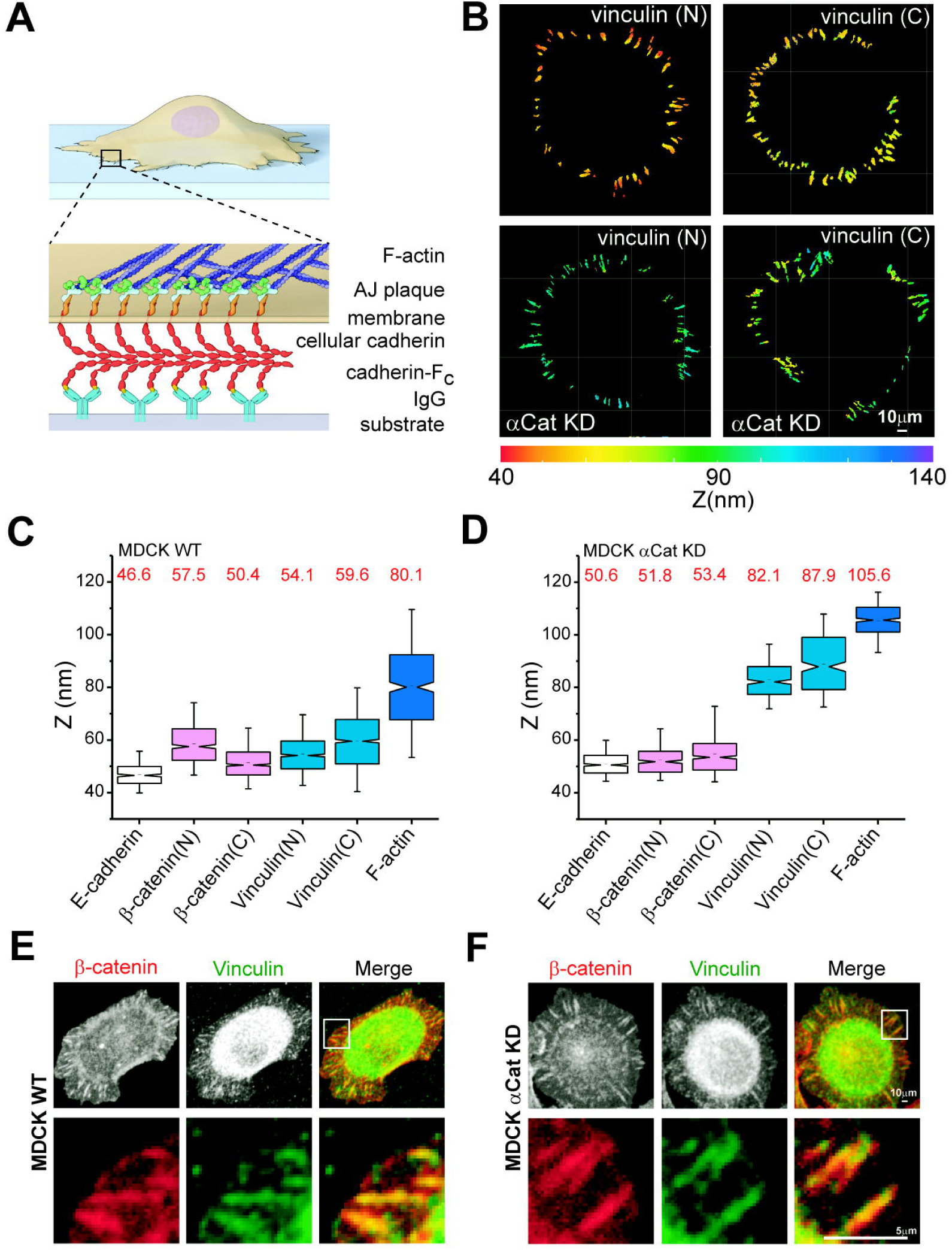
Super-resolution microscopy reveals an alternate nanoscale organization of vinculin in the absence of α-catenin. **A)** Schematic diagram of cadherin biomimetic substrate for high-resolution imaging of cadherin-based adhesions. Imaging substrate (grey) is coated by IgG (blue) and a chimera of F_c_ and E-cadherin extracellular domains (red) to support cell adhesions by cadherin homophilic interactions. **B)** Topographic Map of Protein Z-position. MDCK wt and MDCK α-catenin KD expressing FP probes of indicated proteins were cultured on biomimetic E-cadherin-F_c_ -coated silicon wafer and imaged by surface-generated structured illumination microscopy. Color bar, 40-140 nm. Scale Bar, 10 μm. **C-D)** Alternate z-position of vinculin upon α-catenin depletion. Notched box and whisker plots indicating the median z_centre_ position of indicated proteins in control MDCK (C) and α-catenin KD MDCK (D). Box represents median, 1^st^, and 3^rd^ quartiles; whiskers, 5^th^ and 95^th^ percentiles. **E-F)** Localization of vinculin to cadherin-based adhesions on biomimetic substrate. Immunofluorescence micrographs probing for endogenous β-catenin and vinculin in control MDCK (E) and MDCK αCat KD (F). Bottom panels correspond to insets in top panels. Scale bars, 10 μm (top), 5 μm (bottom).

Next we sought to investigate the alternate localization of vinculin. Since vinculin is able to switch between the compact and the extended conformation, the small z-positional differences between the N-and C- termini observed in MDCK α-catenin KD, probably corresponds to a relatively compact conformation. Interestingly, when we introduced vinculin-T12, a constitutively active vinculin mutant [31] into MDCK α-catenin KD, we observed a highly polarized orientation that effectively span between the cadherin-catenin and actin compartments (Fig. 2A). This indicates that, despite the significant depletion of α-catenin (Fig. 2E), vinculin appears to be able to form a physical connection between the cadherin-catenin compartment and the actin cytoskeleton. We corroborated this observation by probing for the z-position of vinculin head (V_H_) domain, observing a lower z-position proximal to the cadherin-catenin compartment, similar to wt-vinculin in control MDCK (Fig. 1C and 2A). These results thus suggest that once relieved of the autoinhibitory head-tail interaction [32], V_H_ domain of vinculin is capable of interacting with an AJ component, other than α-catenin, within the cadherin-catenin compartment.

**Figure 2.**
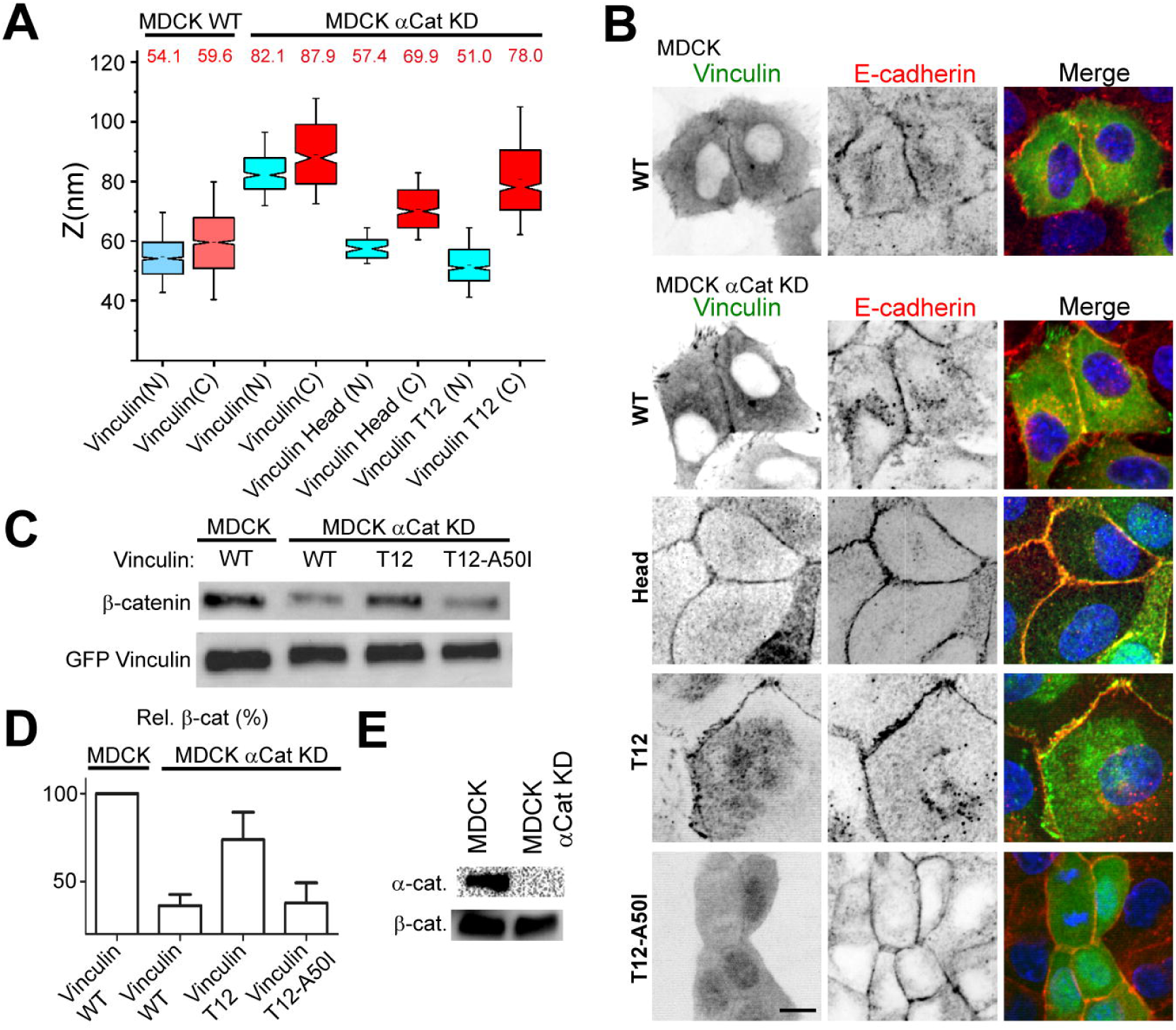
Interaction between activated vinculin and β-catenin. **A)** Activated vinculin was able to span through the core of cadherin-based adhesions. Notched box and whisker plots indicating the median z_centre_ position of vinculin wt and vinculin mutants in MDCK wt and α-catenin KD MDCK. Box represents median, 1^st^, and 3^rd^ quartiles; whiskers, 5^th^ and 95^th^ percentiles. B) Localization of vinculin constructs, as in A, to AJs in MDCK wt and MDCK α-catenin KD in monolayer. Inverted contrast images of vinculin (GFP, green channel) and E-cadherin (antibody probes, red channel), and channel-merged. Scale bar, 10 μm. **C-D)** Co-immunuprecipation of β-catenin with vinculin. Representative western blots (C), probing for β-catenin pulled-down with GFP-vinculin constructs and quantification of β-catenin intensity (D). **E)** Representative western blots probing for Knock-down of α-catenin in MDCK αCat KD cells, in comparison to β-catenin (MDCK wt are used as positive control).

### Interaction between activated vinculin and β-catenin

We observed that the z-position of β-catenin (z_centre_ ~50-60 nm, Fig. 1C-D, Supplementary Table 1) appears to coincide with the N-terminal z-position of vinculin-T12 or vinculin-head (Fig. 2A). These observations raised an interesting possibility of a direct interaction between β-catenin and vinculin, described in a number of earlier studies [25, 33]. We verified this by performing co-immunoprecipitation of vinculin or mutants thereof with β-catenin. In control MDCK, significant co-precipitation was observed between wild-type and β-catenin (Fig. 2C-D). This was significantly diminished to baseline level in MDCK α -catenin KD expressing vinculin-GFP. In contrast, when vinculin-T12-GFP was used for the pull-down, an increase in β-catenin co-precipitation was observed. Since vinculin binding to β-catenin was shown to be abolished by the A50I vinculin mutation [25], while vinculin binding to α-catenin was unperturbed [25], we also made use of the A50I-T12 vinculin mutants. As shown in Fig. 2C-D, this resulted in a diminished β-catenin pull-down. Altogether this data indicated that in the absence of α-catenin, the interaction between wild-type vinculin and β-catenin is minimized. However, the relief of autoinhibition appears to permit vinculin/β-catenin interaction. This is suggestive of the roles of α-catenin in the activation of vinculin to enable vinculin/β-catenin interaction.

### The interaction between β-catenin and activated vinculin support force transmission in AJs

A previous study indicated that the interaction between β-catenin and vinculin stabilizes the cell surface expression of E-cadherin [25]. The N-terminal β-catenin binding site for vinculin was also suggested to be homologous with the amphipathic helices of talin, α-actinin, and α-catenin [25]. As these are well-characterized mechanotransducing binding partners of vinculin [34], we hypothesize that the vinculin/β-catenin interaction may also be mechanically active. To investigate the mechanical roles of the interaction between vinculin and β-catenin, we therefore assessed the tension within the cell-cell junctions of epithelial monolayer by the laser nanoscissor approach. The initial recoil rate following the junction scission has commonly been used as the proxy for tension [17, 35, 36]. As shown in Fig. 3A-C and Supplementary Table 2, in control MDCK monolayer (mEmerald fusion of ZO1 was used as the junction markers), upon laser ablation, the cell-cell junctions underwent rapid ballistic recoil, followed by a more gradual recession. We found that in MDCK α-catenin KD, a significant reduction of the initial recoil rate was registered, indicative of a reduced junction tension relative to control. On the other hand, in MDCK α-catenin KD that expressed vinculin-T12, we observed a rapid junctional recoil with a similar rate to the control. These results suggest that activated vinculin, in the absence of α-catenin, is able to restore high junctional tension, presumably due to the linkage between the actin cytoskeleton and the E-cadherin/β-catenin complex. In contrast, wt-vinculin appears to be impaired in its support of junctional tension, in the absence of α-catenin.

**Figure 3.**
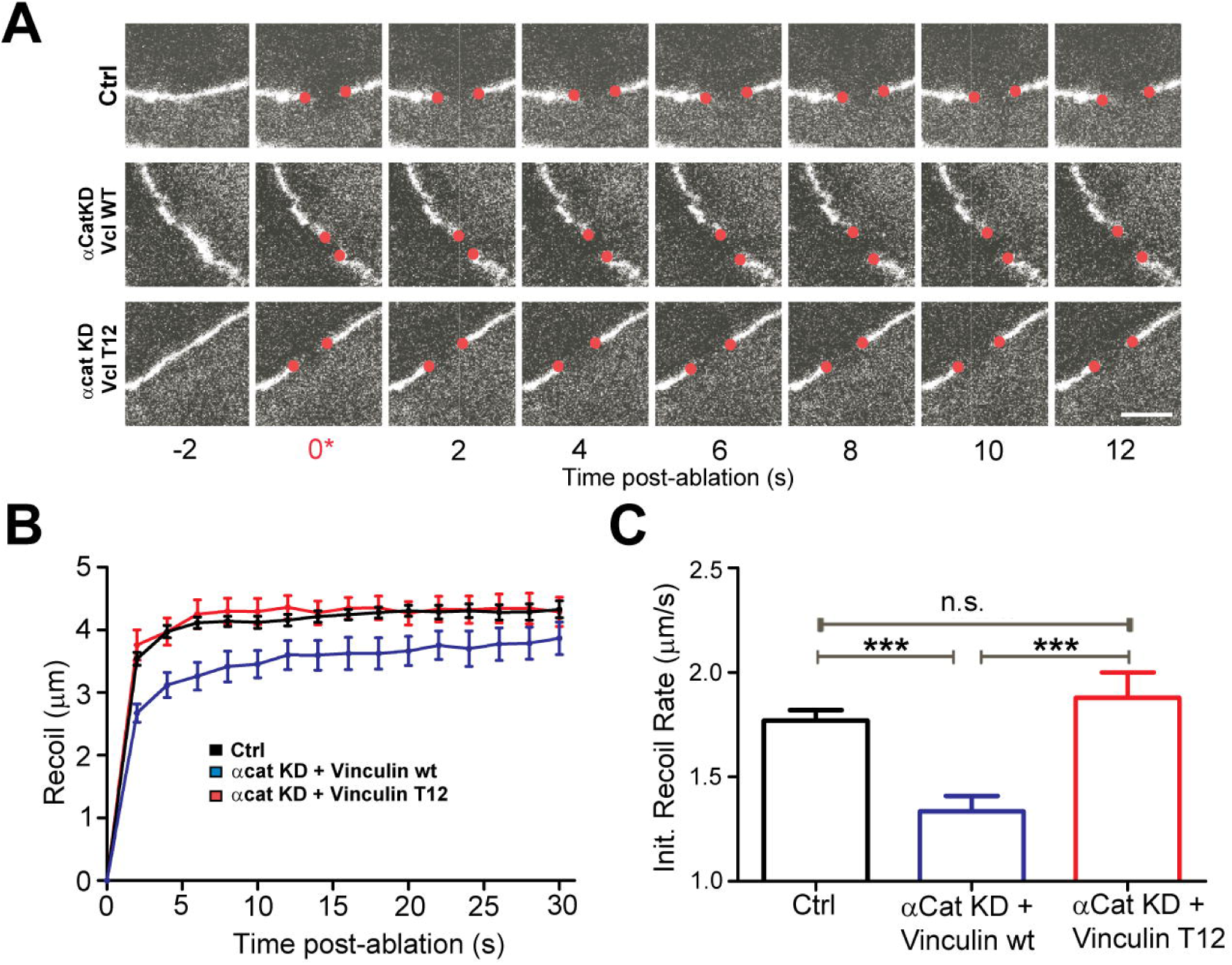
Junctional tension is supported by activated vinculin in the absence of α-catenin. **(A)** Representative time-lapse montages of recoil upon laser ablation. MDCK cells were co-transfected by the indicated constructs as well as ZO1 to mark the cell-cell junctions. The edges of junctions are denoted by red dots. Scale Bar, 10μm. **(B)** Trajectory of junctional recoil following ablation. **C)** Initial recoil rate after laser ablation. Error bars in B-C indicate s.e.m. ***: p < xxx; n.s.: not significant.

### Activated vinculin stabilizes cadherin-catenin complex in AJs

Since the unstructured cytoplasmic domain of E-cadherin serves as the binding site for β-catenin[37], we further investigated the contribution of vinculin/β-catenin interaction to the stability of the cadherin-catenin complexes. We performed Fluorescence Recovery After Photobleaching (FRAP) experiments in MDCK epithelial monolayer, monitoring the intensity of GFP fusion of β-catenin (Fig. 4A). In control MDCK cells, photobleaching was followed by a recovery with a relatively slow half-time of 25.6 s (Fig. 4B an Supplementary Table 3). The extent of recovery is 33 %, indicative of a relatively small dynamically exchanged fraction of β-catenin, consistent with the role of β-catenin in stabilizing E-cadherin in AJs[38]. In MDCK α-catenin KD, we observed a significantly greater recovery fraction, as well as a faster half-time (Fig. 4B-C). These results are suggestive of the roles of α-catenin in both promoting the stability and limiting the mobility of the E-cadherin ternary complex, recently shown to be important for cluster formation[39]. Upon the expression of vinculin T-12, we observed an increase in the immobile fraction, indicative of the stabilization of the junctional cadherin-catenin complex. However, the recovery half-time remains comparable to in the case of MDCK α-catenin KD alone. To evaluate the contribution of the linkage to the actin cytoskeleton for the junctional dynamics, we also made use of the vinculin-head constructs that lack actin binding site (Fig. 4B-D). We observed a relatively high mobile fraction as well as a fast exchange, comparable to the MDCK α-catenin KD. Altogether these results suggest that the direct linkage to the actin cytoskeleton provided by activated vinculin may promote the stability of the cadherin-catenin complexes that are resident in AJs. However, the dynamics of the exchange appears to be insensitive to vinculin activation, presumably since vinculin activation must be preceded by their immobilization in AJs. Here, the absence of α-catenin from the cadherin-catenin complex is likely accounting for the relatively high diffusivity observed.

**Figure 4.**
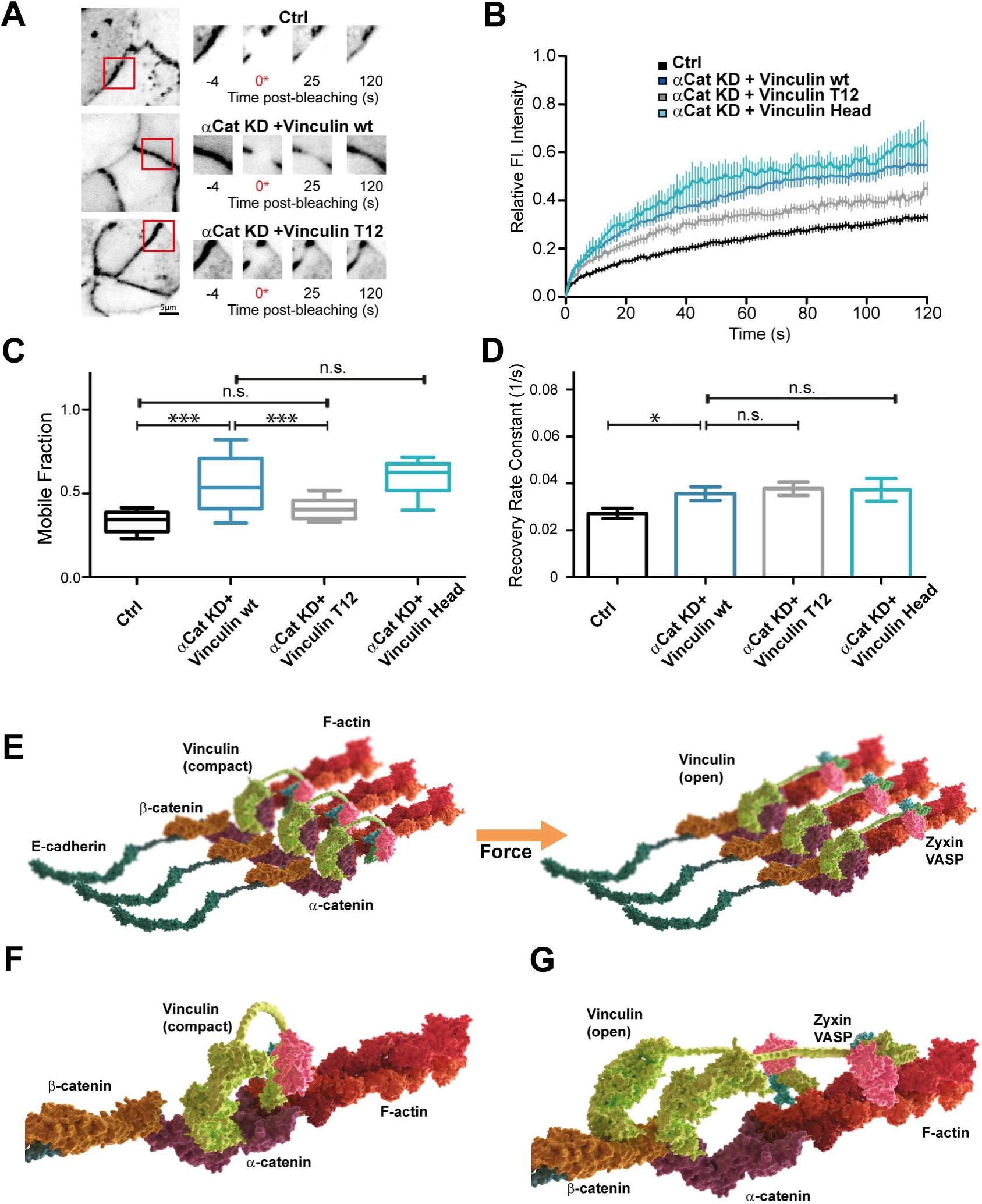
Stabilization of junctional β-catenin by activated vinculin and β-catenin interaction. **A)** Representative time-lapse images from Fluorescence Recovery After Photobleaching (FRAP) of β-catenin. (Right) Montage (zoom 2X) corresponding to red boxes region. Scale bar, 5 μm. **B-D)** FRAP analysis of β-catenin. **B)** Normalized recovery trajectory was fitted to a single exponential model (Supplementary Table 3), with mobile fraction of β-catenin relative to total in (C) and recovery rate constant in **(D)**. **E-G)** Model for two-step connection between cadherin and the actin cytoskeleton. Ternary complex of E-cadherin/β-catenin/α-catenin recruits vinculin via the mechanically gated vinculin binding site on α-catenin **(F)**. **G)** Once activated, vinculin may translocate to form a more direct connection via β-catenin, presumably enabling reinforcement of the cadherin-actin linkage. Error bars indicate s.e.m. *:p<0.05;***: p < 5E-4; n.s.: not significant.

### Conclusion: Implications on Mechanical Force Transmission in AJs

Our data indicated that vinculin could form a direct physical connection between the actin cytoskeleton and β-catenin, bypassing α-catenin. These are reflected in both the organization of vinculin at the nanoscale level, as well as in the regulation of junctional tension in the cell-cell junctions of epithelial monolayer. Such interaction requires vinculin activation, however, which could be achieved in the absence of α-catenin by site-specific mutations. Although based on an α-catenin depletion background, our findings suggest that the nature of the physical linkage between cadherin and actin could be parse-able into distinct steps beyond the current model [15, 16]. First, the ternary complex of E-cadherin/β-catenin/α-catenin may serve as the initial nucleation sites for cell-cell adhesions. Since α-catenin unfolds under a relatively low ~5pN tension [16], this may simultaneously disrupt the autoinhibition and recruit vinculin. Subsequently vinculin may reinforce the connection to the actin cytoskeleton through both the actin-binding C-terminal vinculin-tail (V_T_) domain and probably via the association of the actin elongation factor VASP to the poly-proline-rich linker region [29]. As depicted in Fig. 4E, once activated and fully extended, our results suggest that the V_H_ domain of vinculin could become engaged with the N-terminal site on β-catenin to provide a more direct linkage with the E-cadherin/β-catenin complex. As shown in junctional tension measurements, the activated vinculin/β-catenin interaction could be force-bearing, supporting a comparable magnitude of tension as in the control.

Since α-catenin has been demonstrated to exhibit a catch-bond behaviour with peak lifetime at <10 pN [11], under a situation with higher tension, such as following vinculin recruitment, the bond lifetime is expected to be diminished. We speculate that this may favour the disengagement of the activated vinculin from α-catenin. Since the N-terminal vinculin binding site of β-catenin is in close proximity and does not overlap with α-catenin/β-catenin binding site[40], it is conceivable that vinculin V_H_ domain could translocate to link up with β-catenin directly. This would free up the vinculin binding site on α-catenin, allowing the recruitment and activation of another vinculin molecule. In other words, α-catenin could be considered as a catalyst that activate and potentiate vinculin. Following such activation, the mechanically primed vinculin could translocate to to serve as a mechanical linker, on β-catenin as described in this study, but potentially on other partners such as α-actinin.

We note that an analogous vinculin translocation process has recently been described in integrin-based focal adhesions [41]. Vinculin is first recruited by phosphorylated paxillin into the membrane proximal nanoscale compartment, activated by interaction with talin, and translocated to the upper actin regulatory compartments and the stress fibers. In focal adhesions, vinculin-mediated reinforcement can be readily achieved since there is an abundance of vinculin binding sites, 11 on talin alone [42]. In AJs the number of vinculin binding site is much limited. We speculate that if a comparable molecular design principle is at play, the translocation of activated vinculin to β-catenin could serve as a parallel mechanism in AJs, with the primary site on α-catenin for initial recruitment and activation, and the secondary β-catenin site for reinforcement. Finally, our proposed scheme which bypasses α-catenin may also be consistent with the additional roles of α-catenin such as dimerization and F-actin cross-linking, or interaction with other partners [43]. Within AJs, these processes may occur asynchronously and thus at a given moment, a heterogeneous mix of linkages may be present, which would make it experimentally challenging to dissect their relative contribution to AJ mechanics. Also, to date the interaction between vinculin and β-catenin has been relatively under-studied at the biochemical and biophysical levels. Future experiments would be necessary to fully address the interplay of these connections in regulating AJ properties.

## Supporting information

Combined FRAP_movie

Combined laser ablation_movie

## Acknowledgement

We thank Diego Pitta de Araujo (Mechanobiology Institute) for assistance with graphical model. We gratefully acknowledge funding support from the Ministry of Education (AcRF Tier 2, MOE-T2-1-045 and MOE-T2-1-124 to P.K.; MOE-T2-1-116 to Y.T.), the Mechanobiology Institute seed funding, and the National Research Foundation Competitive Research Programme (NRF2012NRF-CRP001-084).

## Materials and Methods

### Cell culture and sample preparation

Madin Darby Canine Kidney (MDCK) cells and the MDCK cell line with stable expression of α-catenin shRNA were cultured in DMEM media with high glucose, supplemented with 10% Fetal Bovine Serum (FBS), and 100 units/mL of Penicillin/ Streptomycin (Life Technologies, Carlsbad, CA). Both cell lines were kind gifts from W. James Nelson, Stanford University. Cells were transfected with ~10 μg of plasmid DNA encoding the protein of interest by electroporation using Neon^®^ Transfection System (Life Technologies), with the following settings: one 1,650 V pulse with 20 ms width for ~1 × 10^6^ cells. The cell lines used were monthly tested for mycoplasma contamination by PCR method. No cell lines used in this study were found in the database of commonly misidentified cell lines that is maintained by ICLAC and NCBI Biosample. The cell lines were not authenticated.

### Fluorescent Protein Fusion Constructs

FP fusion construct of vinculin and ZO-1 mEmerald were created in the laboratory of M.W. Davidson, The Florida State University, and available from Addgene depository [Addgene #54316]. EGFP-vinculin head (residue 1-258) and the N-terminal fusion of vinculin-T12 were obtained from Addgene (#46270, #46266, contributed by Susan Craig, Johns Hopkins University). The C-terminal fusion of vinculin-T12 and the Y822E and Y822F mutants of vinculins were described previously [17, 44]. All constructs created in-house were verified by sequencing. EGFP-β-catenin expression vector was a gift from B.M. Gumbiner (University of Virginia). Localizations to the cell-cell junctions were verified in MDCK cells plated on fibronectin-coated coverglass prepared by incubating 10 μg / mL of bovine fibronectin (F1141, Sigma) in PBS for 1h at 37 °C in humidified atmosphere, cultured to confluence, and imaged by epifluorescence microscopy.

### Surface modification of imaging substrates by oriented cadherin extracellular domains

Silicon oxide wafers, p-type (100)-orientation with ~500 nm thermal oxide were obtained from a commercial source (Bondatek; Addison Engineering, CA). The thermal oxide thickness was measured using an ellipsometer (UV-VIS-VASE, JA-Woollam) located at SMART (Singapore-MIT alliance for Research and Technology, Singapore). Wafers were then pre-treated as described earlier [17]. Briefly, they were cut into ~ 1.2 cm × 1.2 cm chips with a diamond-tip pen, pre-rinsed with distilled water, sonicated in 100 % Acetone, and then in 1 M potassium hydroxide for 20 min each, followed by a wash in distilled water. Silicon wafers were silanized for protein conjugation by incubation with 3-glycidoxy-propyl-dimethoxy-methylsilane (Sigma) (0.045 % in 100 % EtOH) for 1 h, then cured at 110 ° C for 1 h. Silanized substrates were washed with 70% EtOH first and distilled water, then air-dried by nitrogen. Subsequently, the substrates were incubated with goat anti-human F_c_ fragment specific antibody for E-cadherin substrate (Jackson ImmunoResearch, West Grove, PA), at 1 μg / cm^2^ in 0.1 M pH 8 borate buffer at 4°C overnight in a humidity chamber. After rinsing with PBS, the substrates were neutralized by aminoethoxy-EtOH (Sigma) in NaHCO_3_ (100 mM, pH 8.3) for 1 h. Following a washing step, the substrates were incubated for 2 h with human E-cadherin-F_c_ (R&D system, Minneapolis, MN) at 1 μg / cm^2^, rinsed with PBS, and blocked with 0.2 % pluronic acid (Sigma) in PBS for 20 min at room temperature. All washing steps were done in PBS with Ca^2+^ and Mg^2+^ to avoid perturbing the cadherins. Transfected cells were detached by diluted trypsin (1:5 in PBS) and replated in serum free medium onto the silicon wafers, pre-coated with E-cadherin-F_c_ as described above, and incubated at 37 °C in 5 % CO_2_.

### Nanoscale-precision Z-position measurement by surface-generated structured illumination

Scanning Angle Interference Microscopy (SAIM) was performed as previously described [17]. Briefly, acquisition of raw images was performed on a Nikon Eclipse Ti inverted microscope (Nikon Instruments, Japan), equipped with a motorized TIRF illuminator, using a 60 X NA 1.49 Apo TIRF objective lens, a sCMOS camera (Orca Flash 4.0, Hamamatsu, Japan), and a laser combiner (488 nm and 561 nm, Omicron Laserage, Germany) coupled with a polarization-maintaining optical fiber. For imaging, silicon wafers were placed into a PHEM buffer (without EGTA)-filled 27-mm glass-bottom dish (Iwaki, Japan), with the sample side facing downward, and maintained at neutral buoyancy by a thumb screw. Fluorescence images were acquired, as described previously, between 0° (normal) and 52° with 4° incidence angle interval, using pre-tabulated values for either 488 nm (for eGFP) or 561 nm laser (for photo-converted EosFP). As needed, photoconversion of tdEos2 was carried out using LED excitation and DAPI filter set (Lumencor SOLA, Beaverton, OR). Analysis was performed using a custom written IDL-based software. Binary masks for region of interest (ROI) to analyze were defined by simple thresholding of background subtracted images. Topographic height (z) and other fit parameters were determined by fitting the theoretical angle-dependence curve to experimental angle-dependence intensity by the Levenberg–Marquadt nonlinear least square method, for each pixel in the ROI. The median z (height)-position value of each ROI was denoted z_center_, and used as the representative z-position for a given ROI. Topographic Z map were plotted using color to encode z-position.

### Co-Immunuprecipitation Analysis

Cell lysis, protein extraction and co-IP have been performed following the protocols published earlier[45] with slight modification. MDCK cells and MDCK α-catenin KD cells transfected with different vinculin constructs (eGFP vinculin wt, vinculin T12 eGFP, vinculin T12 A50I eGFP, vinculin head eGFP, vinculin Y822E eGFP and Y822F eGFP) were washed twice in HS buffer (20 mM HEPES pH 7.4, and 150 mM NaCl), and lysed on ice in GFP immunoprecipitation buffer (50 mM Tris-HCl pH 7.6, 150 mM NaCl, 1% NP-40, 0.5% deoxycholate, 100 μl/ml HALT™ Protease/ Phosphatase inhibitor cocktail EDTA-free, 1 mM PMSF). GFP was immuno-precipitated with GFP-Trap_A (ChromoTek) with one-third of the lysates kept for WCL (whole cell lysates) analysis and quantification of total protein levels. Immuno-precipitates were washed four times in GFP-immuno-precipitation buffer. As described earlier [17], SDS-PAGE was then performed using 4–15% Mini-PROTEAN^®^ TGX™ Precast Protein Gels (Bio-Rad), with 30 μg of proteins per band, and subsequently transferred to Immobilon-P PVDF (poly-vinylidene fluoride) membranes. PVDF membranes were blocked with 5% non-fat dry milk (wt/vol) in TBS-T buffer (0.1% vol/vol of Tween-20 in TBS buffer solution, #1706435 Bio-Rad) for 1 h at room temperature, and incubated for 1 h at room temperature or overnight at 4 °C with antibodies in 3% non-fat dry milk (wt/vol) in TBS-T buffer. Antibodies used were: GFP (ab290, Abcam) 1:2000; β -catenin (610154, BD Transduction Laboratories) 1:2000. After primary antibody incubation, PVDF membranes were washed in TBS-T, 2.5% non-fat dry milk (wt/vol) in TBS-T buffer, and 5% non-fat dry milk (wt/vol) in TBS-T buffer (10 min each) consecutively. After these three washings, the membranes were incubated with the appropriated horse radish peroxidase-conjugated antibodies in 3% non-fat dry milk (wt/vol) in TBS-T buffer at a dilution of 1:5000. After secondary antibodies incubation, membranes were washed in TBS-T (3 × 10 min) and then incubated for 3-5 minutes with SuperSignal™ West Pico Chemi-luminescent Substrate. Membranes were imaged with Chemidoc Touch System (Bio-Rad). In case of re-probing by different antibodies, PVDF membranes were stripped using Restore Western Blot Stripping Buffer (21059, Thermo Scientific) for 15 min, washed in TBS-T for 10 min, and blocked in 5% non-fat dry milk (wt/vol) in TBS-T buffer at 4 °C overnight or for 1 h at room temperature.

### Laser ablation of cell-cell junctions

For the measurements of cell-cell junction recoil upon laser ablation, MDCK cells were cultured on fibronectin-coated coverglass prepared as described above. Cells were transfected with the FP fusion expression vectors for ZO-1 as marker for the cell contacts, in conjunction with vinculin constructs as appropriate. UV laser nanoscissor ablation of cell-cell junctions in MDCK epithelial monolayer was performed on a Nikon A1R MP laser scanning confocal microscopy, equipped with an ultraviolet laser (PowerChip PNV-0150-100, Team Photonics: 355 nm, 300 ps pulse duration, 1 kHz repetition rate), as described previously [17, 35]. To allow for simultaneous ablation and live imaging, the UV laser beam, controlled independently from the microscope, was combined onto the optical axis via a customized optical path and a dichroic filter. A custom ImageJ plug-in was used to control both the actuators and the shutter, enabling line ablation of 2-3 μm across the cells by motorized mirror motion. Laser ablation was carried out at the z-plane of adherens junction using 15 nW laser power at the back aperture of the objective lens with an exposure time of 350 ms, controlled by a mechanical shutter (VS25S2ZM0, Uniblitz). Images were acquired every 2 s, starting from 3 frames prior to the ablation until 30 frames post-ablation with a scan speed of 1 frame/sec, and a pinhole size of 74 μm. Image analysis of the recoil speed was performed using MTrackJ plugin in ImageJ by tracking the coordinates of the two edges of the cut in subsequent frames. The recoil speed (μm/sec) was defined as the rate of change of the position of the cut edges. The initial recoil speeds were measured from the first 2 s of recoil after ablation.

### Fluorescence recovery after photobleaching (FRAP) measurements

For the measurements of β-catenin mobility, MDCK cells were cultured until confluent on fibronectin-coated coverglass prepared by incubating 10 μg / mL of bovine fibronectin (F1141, Sigma) in PBS for 1 h at 37 °C in humidified atmosphere. Cells were transfected with the expression vectors for β-catenin fused with eGFP, in conjunction with vinculin constructs tagged with mApple or mcherry tag. Imaging was performed using a Nikon Eclipse Ti-E inverted microscope (Nikon, Japan) equipped with a spinning disk confocal unit, CSU-W1 (Yokogawa, Japan), an iLas^2^ illumination system (Roper Scientific), and a ProEM HS EMCCD camera (Princeton instruments, USA). The objective lens used was a CFI Plan Apo VC 60X NA 1.20 water immersion (Nikon).

Ninety consecutive frames of fluorescence images with 488 nm laser were acquired at 1 s interval. After the fourth acquisition, photobleaching in a circular region-of-interest (ROI) (15 px diameter) was carried out by scanning the 488 nm laser at 100% power for 2 ms. For the analysis, the acquired images were background subtracted and corrected for photobleaching by double normalization method, as follow:

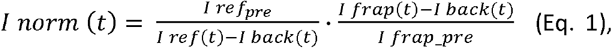

where *I*_*norm*_*(t)* is the normalized intensity; *I*_*frap*_ *(t)* the measured average intensity inside the bleached spot; *I*_*ref*_ *(t)* the measured average reference (an unbleached adhesion) intensity; *I*_*back*_ *(t)* the measured average background intensity outside the cell; subscript _pre indicates the averaged background-substracted intensity in the corresponding ROI before bleaching. For the full scale calibration we used the following formula:

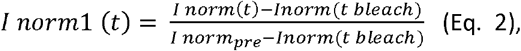

where t_bleach_ is the bleach time. The normalized values were fitted with single exponential function, *I*(*t*) = *P*(1 − *e*^−*kt*^), where P is the mobile fraction and k is the rate constant. The immobile fraction was computed as 1 − *P*. GraphPad Prism 6.0 was used for curve fitting.

## Code Availability

Computation code used in this study is available from the corresponding author upon request.

## Data Availability

All data supporting the conclusions are available from the corresponding author upon reasonable request.

## References

1. Martin, A.C., and Goldstein, B. (2014). Apical constriction: themes and variations on a cellular mechanism driving morphogenesis. Development 141, 1987–1998.

2. Lecuit, T., and Yap, A.S. (2015). E-cadherin junctions as active mechanical integrators in tissue dynamics. Nature cell biology 17, 533–539.

3. Ladoux, B., and Mege, R.M. (2017). Mechanobiology of collective cell behaviours. Nature reviews. Molecular cell biology 18, 743–757.

4. Guo, Z., Neilson, L.J., Zhong, H., Murray, P.S., Zanivan, S., and Zaidel-Bar, R. (2014). E-cadherin interactome complexity and robustness resolved by quantitative proteomics. Science signaling 7, rs7.

5. Leckband, D., and de Rooij, J. (2014). Cadherin adhesion and mechanotransduction. Annual review of cell and developmental biology 30, 291–315.

6. Takeichi, M. (2014). Dynamic contacts: rearranging adherens junctions to drive epithelial remodelling. Nature Reviews Molecular Cell Biology 15, 397–410.

7. Yonemura, S. (2011). Cadherin–actin interactions at adherens junctions. Current opinion in cell biology 23, 515–522.

8. Zaidel-Bar, R. (2013). Cadherin adhesome at a glance. Journal of cell science 126, 373–378.

9. Ladoux, B., Nelson, W.J., Yan, J., and Mege, R.M. (2015). The mechanotransduction machinery at work at adherens junctions. Integrative biology : quantitative biosciences from nano to macro 7, 1109–1119.

10. Xia, S., and Kanchanawong, P. (2017). Nanoscale mechanobiology of cell adhesions. Seminars in cell & developmental biology 71, 53–67.

11. Buckley, C.D., Tan, J., Anderson, K.L., Hanein, D., Volkmann, N., Weis, W.I., Nelson, W.J., and Dunn, A.R. (2014). Cell adhesion. The minimal cadherin-catenin complex binds to actin filaments under force. Science (New York, N.Y 346, 1254211.

12. Borghi, N., Sorokina, M., Shcherbakova, O.G., Weis, W.I., Pruitt, B.L., Nelson, W.J., and Dunn, A.R. (2012). E-cadherin is under constitutive actomyosin-generated tension that is increased at cell-cell contacts upon externally applied stretch. Proceedings of the National Academy of Sciences of the United States of America 109, 12568–12573.

13. Acharya, B.R., and Yap, A.S. (2016). Pli Selon Pli: Mechanochemical Feedback and the Morphogenetic Role of Contractility at Cadherin Cell-Cell Junctions. Current topics in developmental biology 117, 631–646.

14. Kim, T.J., Zheng, S., Sun, J., Muhamed, I., Wu, J., Lei, L., Kong, X., Leckband, D.E., and Wang, Y. (2015). Dynamic visualization of alpha-catenin reveals rapid, reversible conformation switching between tension states. Current biology : CB 25, 218–224.

15. Yonemura, S., Wada, Y., Watanabe, T., Nagafuchi, A., and Shibata, M. (2010). alpha-Catenin as a tension transducer that induces adherens junction development. Nature cell biology 12, 533–542.

16. Yao, M., Qiu, W., Liu, R., Efremov, A.K., Cong, P., Seddiki, R., Payre, M., Lim, C.T., Ladoux, B., Mege, R.M., et al. (2014). Force-dependent conformational switch of alpha-catenin controls vinculin binding. Nature communications 5, 4525.

17. Bertocchi, C., Wang, Y., Ravasio, A., Hara, Y., Wu, Y., Sailov, T., Baird, M.A., Davidson, M.W., Zaidel-Bar, R., Toyama, Y., et al. (2017). Nanoscale architecture of cadherin-based cell adhesions. Nature cell biology 19, 28–37.

18. Barry, A.K., Tabdili, H., Muhamed, I., Wu, J., Shashikanth, N., Gomez, G.A., Yap, A.S., Gottardi, C.J., de Rooij, J., Wang, N., et al. (2014). alpha-catenin cytomechanics--role in cadherin-dependent adhesion and mechanotransduction. Journal of cell science 127, 1779–1791.

19. Valenta, T., Hausmann, G., and Basler, K. (2012). The many faces and functions of beta-catenin. The EMBO journal 31, 2714–2736.

20. Mangold, S., Norwood, S.J., Yap, A.S., and Collins, B.M. (2012). The juxtamembrane domain of the E-cadherin cytoplasmic tail contributes to its interaction with Myosin VI. Bioarchitecture 2, 185–188.

21. Abe, K., and Takeichi, M. (2008). EPLIN mediates linkage of the cadherin catenin complex to F-actin and stabilizes the circumferential actin belt. Proceedings of the National Academy of Sciences of the United States of America 105, 13–19.

22. Knudsen, K.A., Soler, A.P., Johnson, K.R., and Wheelock, M.J. (1995). Interaction of alpha-actinin with the cadherin/catenin cell-cell adhesion complex via alpha-catenin. The Journal of cell biology 130, 67–77.

23. Pokutta, S., Drees, F., Takai, Y., Nelson, W.J., and Weis, W.I. (2002). Biochemical and structural definition of the l-afadin- and actin-binding sites of alpha-catenin. The Journal of biological chemistry 277, 18868–18874.

24. Tachibana, K., Nakanishi, H., Mandai, K., Ozaki, K., Ikeda, W., Yamamoto, Y., Nagafuchi, A., Tsukita, S., and Takai, Y. (2000). Two cell adhesion molecules, nectin and cadherin, interact through their cytoplasmic domain-associated proteins. The Journal of cell biology 150, 1161–1176.

25. Peng, X., Cuff, L.E., Lawton, C.D., and DeMali, K.A. (2010). Vinculin regulates cell-surface E-cadherin expression by binding to beta-catenin. Journal of cell science 123, 567–577.

26. Paszek, M.J., DuFort, C.C., Rubashkin, M.G., Davidson, M.W., Thorn, K.S., Liphardt, J.T., and Weaver, V.M. (2012). Scanning angle interference microscopy reveals cell dynamics at the nanoscale. Nature methods 9, 825–827.

27. Ajo-Franklin, C.M., Ganesan, P.V., and Boxer, S.G. (2005). Variable incidence angle fluorescence interference contrast microscopy for z-imaging single objects. Biophysical journal 89, 2759–2769.

28. Benjamin, J.M., Kwiatkowski, A.V., Yang, C., Korobova, F., Pokutta, S., Svitkina, T., Weis, W.I., and Nelson, W.J. (2010). αE-catenin regulates actin dynamics independently of cadherin-mediated cell–cell adhesion. The Journal of cell biology 189, 339–352.

29. Leerberg, J.M., Gomez, G.A., Verma, S., Moussa, E.J., Wu, S.K., Priya, R., Hoffman, B.D., Grashoff, C., Schwartz, M.A., and Yap, A.S. (2014). Tension-sensitive actin assembly supports contractility at the epithelial zonula adherens. Current biology : CB 24, 1689–1699.

30. Bois, P.R., Borgon, R.A., Vonrhein, C., and Izard, T. (2005). Structural dynamics of alpha-actinin-vinculin interactions. Molecular and cellular biology 25, 6112–6122.

31. Cohen, D.M., Kutscher, B., Chen, H., Murphy, D.B., and Craig, S.W. (2006). A conformational switch in vinculin drives formation and dynamics of a talin-vinculin complex at focal adhesions. The Journal of biological chemistry 281, 16006–16015.

32. Cohen, D.M., Chen, H., Johnson, R.P., Choudhury, B., and Craig, S.W. (2005). Two distinct head-tail interfaces cooperate to suppress activation of vinculin by talin. The Journal of biological chemistry 280, 17109–17117.

33. Hazan, R.B., Kang, L., Roe, S., Borgen, P.I., and Rimm, D.L. (1997). Vinculin is associated with the E-cadherin adhesion complex. The Journal of biological chemistry 272, 32448–32453.

34. Peng, X., Nelson, E.S., Maiers, J.L., and DeMali, K.A. (2011). New insights into vinculin function and regulation. Int Rev Cell Mol Biol 287, 191–231.

35. Ravasio, A., Cheddadi, I., Chen, T., Pereira, T., Ong, H.T., Bertocchi, C., Brugues, A., Jacinto, A., Kabla, A.J., Toyama, Y., et al. (2015). Gap geometry dictates epithelial closure efficiency. Nature communications 6, 7683.

36. Hara, Y., Shagirov, M., and Toyama, Y. (2016). Cell boundary elongation by non-autonomous contractility in cell oscillation. Current Biology 26, 2388–2396.

37. Huber, A.H., and Weis, W.I. (2001). The structure of the β-catenin/E-cadherin complex and the molecular basis of diverse ligand recognition by β-catenin. Cell 105, 391–402.

38. Orsulic, S., Huber, O., Aberle, H., Arnold, S., and Kemler, R. (1999). E-cadherin binding prevents beta-catenin nuclear localization and beta-catenin/LEF-1-mediated transactivation. Journal of cell science 112, 1237–1245.

39. Biswas, K.H., Hartman, K.L., Zaidel-Bar, R., and Groves, J.T. (2016). Sustained alpha-catenin Activation at E-cadherin Junctions in the Absence of Mechanical Force. Biophysical journal 111, 1044–1052.

40. Klionsky, D.J., Abdelmohsen, K., Abe, A., Abedin, M.J., Abeliovich, H., Acevedo Arozena, A., Adachi, H., Adams, C.M., Adams, P.D., Adeli, K., et al. (2016). Guidelines for the use and interpretation of assays for monitoring autophagy (3rd edition). Autophagy 12, 1–222.

41. Case, L.B., Baird, M.A., Shtengel, G., Campbell, S.L., Hess, H.F., Davidson, M.W., and Waterman, C.M. (2015). Molecular mechanism of vinculin activation and nanoscale spatial organization in focal adhesions. Nature cell biology 17, 880–892.

42. Critchley, D.R. (2009). Biochemical and structural properties of the integrin-associated cytoskeletal protein talin. Annu Rev Biophys 38, 235–254.

43. Drees, F., Pokutta, S., Yamada, S., Nelson, W.J., and Weis, W.I. (2005). Alpha-catenin is a molecular switch that binds E-cadherin-beta-catenin and regulates actin-filament assembly. Cell 123, 903–915.

44. Bays, J.L., Peng, X., Tolbert, C.E., Guilluy, C., Angell, A.E., Pan, Y., Superfine, R., Burridge, K., and DeMali, K.A. (2014). Vinculin phosphorylation differentially regulates mechanotransduction at cell-cell and cell-matrix adhesions. The Journal of cell biology 205, 251–263.

45. Bays, J.L., Peng, X., Tolbert, C.E., Guilluy, C., Angell, A.E., Pan, Y., Superfine, R., Burridge, K., and DeMali, K.A. (2014). Vinculin phosphorylation differentially regulates mechanotransduction at cell–cell and cell–matrix adhesions. The Journal of cell biology 205, 251–263.

