## Supplementary tables for "Mechanical Roles of Vinculin/β-catenin interaction in Adherens Junction"

**Supplementary Table 1:** Statistics of Protein Z-position as measured by surface-generated structured illumination microscopy. The position of fluorophores at the N- or C- termini are indicated.

| Probes | cell type | z_centre_ (median, nm) | N_Adh_ | N_cells_ | Mean (nm) | Std.dev (nm) |
| --- | --- | --- | --- | --- | --- | --- |
| E-Cadherin (C) | MDCK | 46.56 | 352 | 20 | 46.95 | 4.72 |
| β-catenin (N) | MDCK | 57.49 | 605 | 39 | 58.61 | 8.18 |
| β-catenin (C) | MDCK | 50.45 | 898 | 22 | 51.36 | 6.72 |
| Vinculin (N) | MDCK | 54.13 | 1079 | 42 | 54.73 | 7.97 |
| Vinculin (C) | MDCK | 59.56 | 1426 | 28 | 59.55 | 11.81 |
| F-Actin | MDCK | 80.1 | 435 | 23 | 80.57 | 16.86 |
| E-Cadherin (C) | MDCK αcat KD | 50.57 | 478 | 14 | 51.04 | 4.61 |
| β-catenin (N) | MDCK αcat KD | 51.78 | 385 | 15 | 52.4 | 5.72 |
| β-catenin (C) | MDCK αcat KD | 53.43 | 632 | 22 | 54.73 | 8.21 |
| Vinculin (N) | MDCK αcat KD | 82.11 | 324 | 12 | 82.99 | 7.37 |
| Vinculin (C) | MDCK αcat KD | 87.86 | 295 | 11 | 88.88 | 11.24 |
| F-Actin | MDCK αcat KD | 105.57 | 397 | 10 | 105.37 | 4.3 |
| Vinculin head (N) | MDCK αcat KD | 57.42 | 85 | 6 | 57.52 | 3.81 |
| Vinculin TL-TS | MDCK αcat KD | 69.94 | 346 | 14 | 70.68 | 7.31 |
| Vinculin-T12 (N) | MDCK αcat KD | 50.98 | 528 | 17 | 51.91 | 7.2 |
| Vinculin-T12 (C) | MDCK αcat KD | 78.01 | 582 | 15 | 80.66 | 13.31 |

**Supplementary Table 2:** Statistics of cell-cell junction tension measurements by Laser nanoscissor (Fig. 3).

|  | n | Mean Recoil Rate (μm/s) | Std. dev. (μm/s) | s.e.m (μm/s) |
| --- | --- | --- | --- | --- |
| MDCK ctrl | 15 | 1.77 | 0.198 | 0.0511 |
| MDCK αcat KD | 13 | 1.34 | 0.264 | 0.0733 |
| MDCK αcat KD + T12 | 14 | 1.88 | 0.451 | 0.121 |

**Supplementary Table 3:** Statistics of FRAP measurements (Fig. 4). Fit of FRAP curves are fitted to a single exponential function, *I(t) = p(1-e^-kt^)*. Half-time is *ln2/k*.

|  | n | p, mobility fraction | | | k (s^-1^) | | | t_1/2_ (s) |
| --- | --- | --- | --- | --- | --- | --- | --- | --- |
|  |  | Mean | Std. dev. | s.e.m. | Mean | Std. dev. | s.e.m. | Mean |
| MDCK ctrl | 26 | 0.331 | 0.0676 | 0.0133 | 0.0271 | 0.0111 | 0.0022 | 25.6 |
| MDCK αCat KD | 31 | 0.553 | 0.177 | 0.0318 | 0.0356 | 0.0163 | 0.0029 | 19.5 |
| MDCK αCat KD + Vinculin-T12 | 19 | 0.402 | 0.0718 | 0.0165 | 0.0377 | 0.0121 | 0.0029 | 18.4 |
| MDCK αCat KD + Vinculin- head | 6 | 0.598 | 0.112 | 0.0455 | 0.0372 | 0.0121 | 0.0049 | 18.6 |
